# SELECT 2.0: Refined and open access SELection Endpoints in Communities of bacTeria (SELECT) method to determine concentrations of antibiotics that may select for antimicrobial resistance in the environment

**DOI:** 10.64898/2026.03.30.713945

**Authors:** April Hayes, Suzanne Kay, Christopher Lowe, William Gaze, Mario Recker, Angus Buckling, Aimee K. Murray

## Abstract

Antimicrobial resistance (AMR) is a significant threat to human and animal health, the global economy, and food security. The One Health approach to AMR recognises the role of the environment in the evolution, emergence, and dissemination of AMR. This is partly due to anthropogenic pollution that releases AMR organisms alongside cocktails of compounds that may select for AMR *in situ*, which then pose an exposure risk to humans and animals. This has spurred growing interest from cross-sectoral stakeholders in environmental risk assessment of antibiotics, with regards to their selective potential. Many different approaches have been used to determine the lowest concentration of an antibiotic that may select for AMR. Debates continue regarding which individual approach, if any, may be best for determining selective concentrations of antibiotics for environmental risk assessment purposes. This paper contributes to this discourse by refining a previously published method ‘SELECT’ to generate predicted no effect concentrations for resistance (PNECRs) for 32 antibiotics, with open access code and walk-through. The PNECRs presented here is the largest number generated using a single experimental approach that will aid future efforts towards creating a standardised test.

## 1.1 Introduction

In 2019, approximately five million deaths were associated with antibiotic resistance [1], and antimicrobial resistance (AMR) is predicted to become the leading cause of death worldwide by 2050 [2]. AMR is a quintessential One Health issue [3], spanning human, animal, and environmental sectors. Internationally, there is growing concern as to whether antibiotic pollution can select for AMR in the environment and subsequently pose a risk to human health [4, 5]. Previous research (including but not limited to [6–8]) has shown that low, ng/L-µg/L environmentally relevant concentrations of antibiotics can select for AMR. However, current environmental risk assessment approaches rely on established ecotoxicological tests to determine whether compounds pose a risk [9], and these do not directly measure selection for AMR. Previous comparisons have also demonstrated that these ecotoxicological tests can sometimes generate effect concentration data that are higher than the concentrations shown to select for AMR [10, 11], raising concerns that environmental risk assessments based solely on ecotoxicological endpoints may overlook AMR selection risk.

The minimal selective concentration (MSC) is the lowest concentration of antibiotic that selects for resistance and was originally determined in single species competition experiments where isogenic resistant and susceptible bacterial strains were co-cultured at a range of antibiotic concentrations [6]. Additional research has built upon this approach by using complex communities of bacteria and tracking enrichment of resistant strains and/or genes using a combination of culture based and molecular techniques, such as qPCR, amplicon, and metagenomic sequencing (for examples, see [7, 12, 13]). Additional insights into the dynamics and features of sub-inhibitory AMR selection have also been gained through such experiments, including the MSC being increased in the presence of a community [14], the introduction of the concept of the minimum increased persistence concentration [13], and that sub-inhibitory concentrations can lead to rapid resistance evolution [15, 16]. However, the heterogeneity of experimental systems, model organism(s), experimental conditions, and endpoints measured (e.g., specific antibiotic resistance genes) renders comparison of data generated across studies difficult [17]. Furthermore, as this is an emerging research area, the amount of data available remains limited, with several antibiotics being preferentially studied across experimental systems, rather than a range of antibiotics studied using the same experimental approach [17, 18].

In addition to these experimental approaches, several attempts have been made to predict antibiotic MSCs using single-species minimum inhibitory concentration data [19–23]. The most notable of these [19] has gained traction internationally, with generated values being included in WHO Guidance [24] and partially adopted by the pharmaceutical industry as part of their AMR Industry Alliance Antibiotic Manufacturing Standard [25]. However, these data may not accurately reflect AMR selection within complex microbiomes in which most bacteria live, with evidence demonstrating that the presence of a microbial community alters selection for AMR [14, 26]. The pros and cons of different approaches have been discussed elsewhere [17], but it is mutually agreed that values should be updated and reviewed regularly as new data emerge [24], whilst keeping human and environmental health protection at the forefront of any decision making.

Predicted no effect concentrations for resistance (PNECRs) derived from minimum inhibitory concentration (MIC) data or other growth-based data are attractive given the ability to rapidly generate large PNECR datasets using a single, cost-effective approach. Previously, a low-cost, rapid experimental method to determine PNECRs of antibiotics called ‘SELECT’ (SELection Endpoints in Communities of bacTeria) was developed [8]. It was based on the observation that reduction in growth rate of susceptible strains is used to determine the MSC [6], and reduction in growth been shown to be the key parameter for predicting the MSC [27]. SELECT uses a complex community of bacteria to attempt to represent the complex interactions (e.g., competitive, cooperative, protective) that occur in bacterial communities. It measures significant reductions in overall community growth resulting from antibiotic exposure. It was previously shown that SELECT may be a suitable method to quickly generate experimental PNECRs with a single experimental approach, as SELECT PNECRs were similar to PNECRs based on selection for a commonly used AMR marker gene, *intI1*, following 7-day, serial passage antibiotic exposure experiments [8]. Therefore, the phenotypic SELECT method may be a reasonable proxy for genotypic selection within the same experimental system, and provides a low-cost, quick approach to rapidly generate predicted effect concentrations when longer (week-long) experiments, with subsequent phenotyping and genotyping (e.g. qPCR for target genes) is not possible (due to cost and time constraints).

[28, 29]In this study, we refined the statistical approach and produced open access, reproducible code to facilitate further studies and use of the SELECT 2.0 method. The previous method relied on binary definitions of statistical significance, and was strongly affected by random error, which is inherent in community experiments. There has been consistent movement towards modelled effect concentration (EC) data in ecotoxicological risk assessments over statistically derived LOECs and NOECs [28, 29]. Some of the limitations of LOECs and NOECs is that they are significantly impacted by spacing of test concentrations (i.e., 2- or 10-fold dilutions), are constrained to only the concentrations tested, and that they are unable to convey the magnitude of impact [29]. Therefore, in SELECT 2.0, we have moved from a LOEC/NOEC to EC based analysis. Refining this method ensures a more rigorous approach to defining effect concentrations, by considering fitted curves and estimated effect concentrations. We then used SELECT 2.0 to generate 32 putative PNECRs for antibiotics spanning all 8 antibiotic classes [30], generating the largest database of experimental PNECRs using a single method to date. The limitations of SELECT 2.0 are discussed, alongside how these may be addressed in future studies, or further refinement of the method. The PNECR data generated can be used to inform ongoing discussions around environmental quality standards for antibiotics and environmental risk assessment of antibiotics, to reduce the risk of AMR selection in the environment, thereby reducing human exposure risks, and improving antibiotic stewardship.

## 2. Methods

### 2.1.1 Antibiotic stocks

All antibiotic stocks were prepared and stored in single use aliquots at −20°C before use, Antibiotics tested were: amoxicillin, ampicillin, azithromycin, cefotaxime, ceftiofur, ceftriaxone, clarithromycin, chloramphenicol, ciprofloxacin, colistin, doripenem, doxycycline, enrofloxacin, erthryomycin, florfenicol, gentamicin, imipenem, kanamycin, meropenem, nitrofurantoin, norfloxacin, ofloxacin, oxytetracycline, penicillin, streptomycin, sulfadiazine, sulfamethoxazole, sulfapyridine, tetracycline, thiamphenicol and vancomycin. All details on solvents, CAS numbers and product codes can be found in Supplementary Table 1.

### 2.1.2 Standard SELECT assay conditions

All SELECT assays were conducted as previously [8]. Untreated sewage samples were collected from a wastewater treatment plant serving c.a. 43,000 people in June 2022. Samples were mixed 1:1 with 40% sterile glycerol before being frozen at −70°C. Thawed samples were spun down at 14.8G for 2 minutes and the pellet resuspended in 0.85% sterile NaCl (Sigma Aldrich CAS:7647-14-5). This was performed twice to minimise nutrient and chemical carry over. Sewage communities were inoculated into sterile Iso-sensitest broth (Oxoid LOT:3717183) at 10% v/v and 200µl transferred into wells in a 96 well plate (Cyto-One). Then, an aliquot of the same sewage-inoculated Iso-sensitest broth was spiked with double the highest antibiotic test concentration and two-fold serially diluted down the plate. No antibiotic and sterile broth controls were also included. All assays had six replicates per treatment. Separate solvent control assays were also conducted as required to ensure solvents did not affect community growth. Solvent data is also deposited at Zenodo.

Filled 96 well plates were then sealed with an optical seal (MicroAmp) and incubated at 37°C in a MultiSkan plate reader with optical density at 600nm (OD600) readings taken every ten minutes for twelve hours. This is an increase in measuring frequency compared to the previous method, which measured OD every hour as this is likely to produce smoother OD curves that improve OD modelling OD. All raw readings for each antibiotic are in the associated data deposited at Zenodo.

The highest antibiotic test concentration was the European Committee on Antimicrobial Susceptibility Testing (EUCAST) clinical breakpoint for Enterobacterales [31], where available. For antibiotics with no such breakpoint, one of three different approaches were taken. 1) The standard veterinary/clinical dose (mg/kg body weight) was calculated and converted into mg/L start concentration (for florfenicol, enrofloxacin, sulfadiazine, sulfapyridine, and vancomycin). 2) The MIC for a similar antibiotic was used (for thiamphenicol, doxycycline, oxytetracycline, amoxicillin). 3) Trial and error was used by repeat assays with increasing start concentration until growth was affected (for penicillin, sulfamethoxazole, and streptomycin). Notably, azithromycin had an EUCAST breakpoint MIC but the results using this were inconclusive, so a higher starting concentration was used.

### 2.1.3 Solvent Control for NaOH

0.5M NaOH was the only solvent used in this study which had not previously been tested to identify effects on bacterial growth; therefore, we carried out a solvent control. Wastewater influent was thawed and treated as above, and 3µL of 0.5M NaOH was added to the 1.5mL broth and bacteria aliquot. Six wells of the 96 well plate were filled with 200µL broth and bacteria, plus 200µL of the NaOH-bacteria spiked broth, mixed, and 200µL removed to leave a final volume of 200µL. Six wells were left with no NaOH addition as a positive control, and six wells contained only broth as a negative control. The plate was sealed and treated as above. Change in OD did not differ from the positive control growth over 12 hours, with confidence intervals overlapping for both the NaOH treated and control communities (data not shown).

### 2.2 Data Analysis

All statistical analyses and visualisations in this study were conducted and produced using R v4.4.0 in R Studio, using *tidyverse* [32] to manipulate dataframes. For the visualisations, *ggplot2* [33], and *MetBrewer* [34] were used.

#### 2.2.1 Original and refined statistical approach

The original SELECT 1.0 method [8] determined a lowest observed effect concentration (LOEC) by finding the hour time point during exponential growth phase with the strongest dose response relationship (using either Pearson’s or Spearman’s Rank tests, depending on normality of data). The LOEC was defined as the test concentration where overall OD600 was significantly reduced, compared to the no antibiotic control using a Dunn’s test. The no observed effect concentration (NOEC) is the concentration directly below the LOEC, which had no significant effect on growth.

The SELECT 2.0 approach instead determines the effect concentration where overall growth (maximum density) during exponential growth phase decreases by 1% (an Effect Concentration of 1%, or EC1), when fitting a dose response curve. A four-parameter log-logistic curve was fitted to each antibiotic treatment using *drc* v3.0-1 [35]. The fit of the model was checked, and estimated effective doses of 1, 5, and 10, with 95% confidence intervals were generated using *drc::ED*. PNECRs are then generated by applying an assessment factor of 10 to the EC1. We have adopted this approach to better align with the European Medicines Agency recommendations that EC10s are preferable to LOECs for assessment of antibiotics, where a reliable dose response curve is present [9]. Furthermore, we argue that an EC1 should be used instead of an EC10 since bacteria grow significantly more quickly than most other organisms, at higher densities, and therefore an EC1 would be the most protective choice for environmental protection. More details on this decision are included in the results and the discussion.

All the code to determine the EC1 from the raw data are deposited at Zenodo and a tutorial available on GitHub here: https://github.com/ahayyes/SELECT2.0_Tutorial.

#### 2.2.2 Comparison across studies

The data generated in this study were compared to data collected in the meta-analysis by Murray et al., [17]: [6–8, 12–14, 20–23, 36–45] and one further study Lakhey et al., [46]. The database generated in the 2024 study was filtered to include only antibiotics, and only included those studies where selection was observed. All datapoints from these studies were then used to compare to PNECRs generated using SELECT 2.0.

## 3 Results

### 3.1 Dataset of PNECRs derived using SELECT 2.0

In this study, we present the SELECT 2.0 as a refined analysis method to identify effect concentrations of antibiotics using reduction in growth as a proxy for selection for AMR. We determined effect concentrations (EC1s) and resultant PNECRs for 32 different antibiotics across 11 classes (Figure 1A, Table 1, Supplementary Table 2). We found that vancomycin and penicillin had the highest PNECRs of 954µg/L and 263µg/L respectively, and the lowest PNECRs were for ceftriaxone (0.00003µg/L) and ciprofloxacin (0.005µg/L). In general, the quinolones and beta-lactam antibiotics had lower PNECRs, and the aminoglycosides and glycopeptides had higher PNECRs (Figure 1). There was greater variation in PNECRs within the quinolone, beta-lactam, sulfonamide, and macrolide classes than in other antibiotic classes, though a few classes contained only one antibiotic (glycopeptides, dihydropyrimidine).

**Figure 1.**
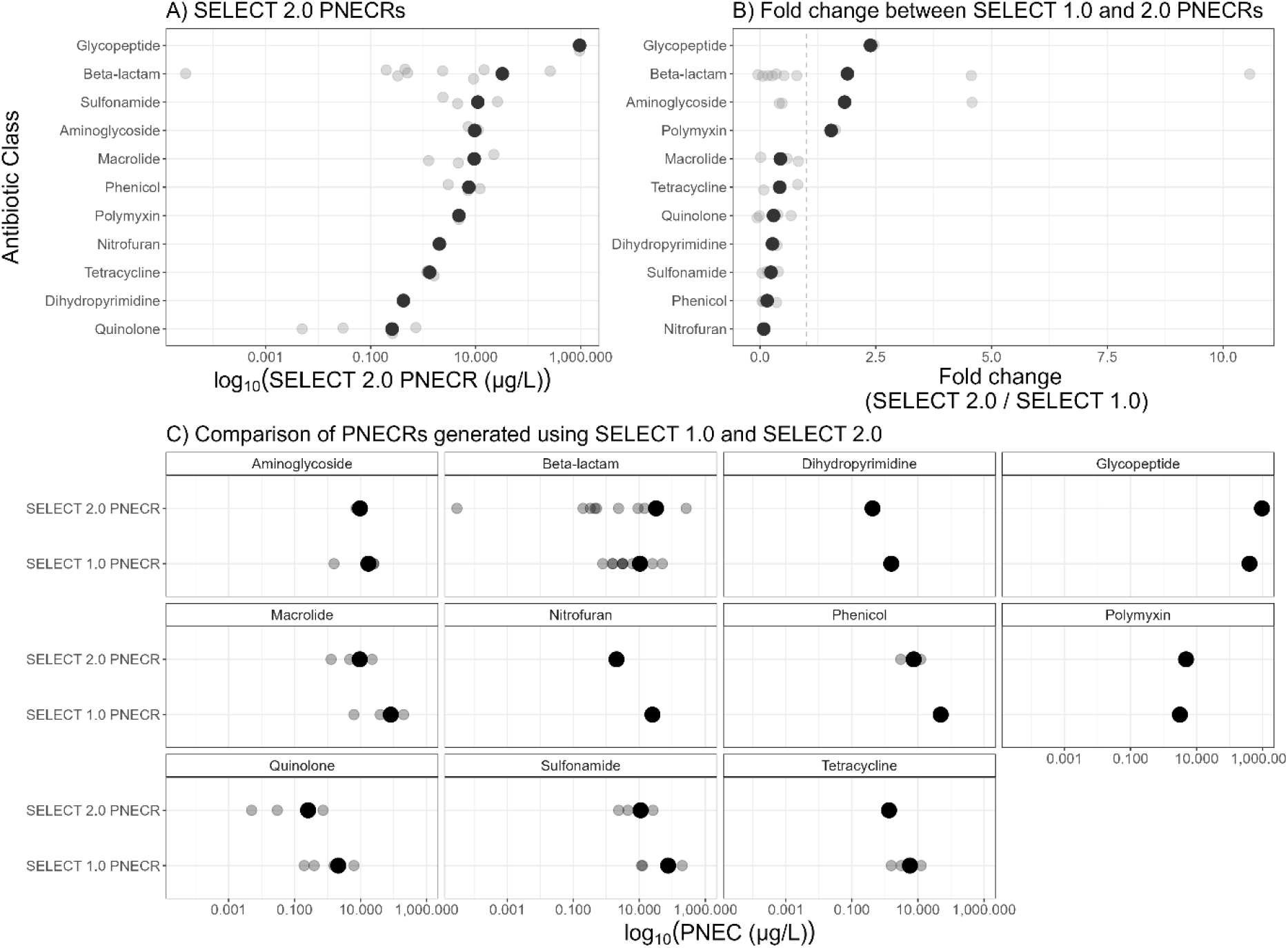
A) PNECRs produced in this study using SELECT 2.0 by antibiotic class. B) Fold change between PNECRs generated by SELECT 1.0 and SELECT 2.0. C) Comparison of PNECRs generated by SELECT 1.0 and 2.0.

**Table 1.** PNECRs generated using the SELECT 1.0 and 2.0 methods, with fold change between the two. All numbers rounded to three significant figures.

| **Antibiotic** | **Class** | **SELECT 1.0 PNECR (µg/L)** | **SELECT 2.0 PNECR (µg/L)** | **Fold change (2.0/1.0)** |
|---|---|---|---|---|
| Gentamicin | Aminoglycoside | 25 | 11.2 | 0.448 |
| Kanamycin | Aminoglycoside | 25 | 10.2 | 0.408 |
| Streptomycin | Aminoglycoside | 1.56 | 7.23 | 4.63 |
| Amoxicilin | Beta-lactam | 3.13 | 14.5 | 4.63 |
| Ampicillin | Beta-lactam | 50 | 9.14 | 0.183 |
| Cefotaxime | Beta-lactam | 1.56 | 0.507 | 0.325 |
| Ceftiofur | Beta-lactam | 3.13 | 2.36 | 0.754 |
| Ceftriaxone | Beta-lactam | 1.56 | 0.0000300 | 0.0000192 |
| Doripenem | Beta-lactam | 0.781 | 0.329 | 0.421 |
| Imipenem | Beta-lactam | 3.13 | 0.451 | 0.144 |
| Meropenem | Beta-lactam | 6.25 | 0.199 | 0.0318 |
| Penicillin | Beta-lactam | 25 | 263 | 10.5 |
| Trimethoprim | Dihydropyrimidine | 1.56 | 0.421 | 0.270 |
| Vancomycin | Glycopeptide | 400 | 954 | 2.39 |
| Azithromycin | Macrolide | 39.1 | 22.2 | 0.568 |
| Clarithromycin | Macrolide | 200 | 1.27 | 0.00635 |
| Erythromycin | Macrolide | 6.25 | 4.68 | 0.749 |
| Nitrofurantoin | Nitrofuran | 25 | 2.04 | 0.0816 |
| Chloramphenicol | Phenicol | 50 | 3.01 | 0.0602 |
| Florfenicol | Phenicol | 46.9 | 12.1 | 0.258 |
| Thiamphenicol | Phenicol | 50 | 7.18 | 0.144 |
| Colistin Sulfate | Polymyxin | 3.13 | 4.8 | 1.53 |
| Ciprofloxacin | Quinolone | 0.195 | 0.00498 | 0.0255 |
| Enrofloxacin | Quinolone | 6.25 | 0.0299 | 0.00478 |
| Norfloxacin | Quinolone | 1.56 | 0.727 | 0.466 |
| Ofloxacin | Quinolone | 0.391 | 0.262 | 0.670 |
| Sulfadiazine | Sulfonamide | 11.7 | 4.53 | 0.387 |
| Sulfamethoxazole | Sulfonamide | 12.5 | 2.38 | 0.190 |
| Sulfapyridine | Sulfonamide | 200 | 26.1 | 0.131 |
| Doxycycline | Tetracycline | 3.13 | 1.18 | 0.377 |
| Oxytetracycline | Tetracycline | 1.56 | 1.2 | 0.769 |
| Tetracycline | Tetracycline | 12.5 | 1.61 | 0.129 |

We calculated standard error and 95% confidence intervals for EC1, EC5, and EC10s generated in this study (Supplementary Figure 1). We found that in general, confidence intervals for EC1s were proportionally smaller than those for EC5s and EC10s, although this varied by antibiotic. Some antibiotics, such as erythromycin, clarithromycin, and colistin had significantly larger confidence intervals around the EC10 estimate compared to the EC1 estimates. We tested whether there were differences between ECXs across all antibiotics, and found that EC1 was significantly lower than EC10 (p=0.013), but that neither EC1 and EC5, nor EC5 and EC10 were significantly different to each other (p=0.12, and p=0.64 respectively). All antibiotic PNECRs were within the experimentally tested concentration range, except ceftriaxone, which had an EC1 that was not experimentally tested (EC1 = 0.0003µg/L, with the lowest tested concentration 0.244µg/L).

### 3.2 Comparison of LOEC and EC1 SELECT PNECRs

We analysed all data generated in this study using the SELECT 1.0 (LOEC/NOEC analysis) and compared it to the 2.0 results (EC analysis). Most of the PNECRs generated using either method were similar across the antibiotic classes, with only a few showing large differences (Table 1, Figure 1B, 1C). All EC1s (version 2.0) and LOECs (version 1.0) generated are present in Supplementary Table 2. Most PNECRs generated using the SELECT 2.0 method were more protective than those generated using SELECT 1.0. Most fold change differences between the two methods were small, with most antibiotics having fold change between 0.5-1. However, ceftriaxone had a fold change of 0.0000192, clarithromycin had a fold change of 0.00635, and penicillin had a fold change of 10.5. When averaging across all antibiotics, there was a fold change difference of 0.99 (95% confidence interval ±0.75) between SELECT 2.0 and 1.0. All class level fold changes can be found in Table 2.

**Table 2.** Mean fold changes per antibiotic class, with standard deviation and 95% confidence interval. Dihydropyrimidine, glycopeptide, nitrofuran, and polymyxin only included one antibiotic within the class, therefore means could not be generated.

| **Class** | **Mean fold change** | **Standard deviation** | **95% confidence interval** |
|---|---|---|---|
| Aminoglycoside | 1.83 | 2.42 | 6.02 |
| Beta-lactam | 1.89 | 3.55 | 2.73 |
| Macrolide | 0.441 | 0.387 | 0.962 |
| Phenicol | 0.154 | 0.0992 | 0.247 |
| Quinolone | 0.292 | 0.330 | 0.525 |
| Sulfonamide | 0.236 | 0.134 | 0.333 |
| Tetracycline | 0.426 | 0.323 | 0.804 |
| **TOTAL**<br>***(averaged across all classes)*** | **0.990** | **2.08** | **0.749** |

### 3.2 Comparison with data from other studies

We collected PNECR data two previous works to compare results produced in this study with those from other studies investigating PNECRs across a range of experimental systems. PNECRs generated using the SELECT 2.0 method are lower or on the lower end of the range of those generated previously (Figure 2, Supplementary Table 4). When averaged within class, almost all SELECT 2.0 PNECRs are lower than those generated from data in other studies (Supplementary Table 5), with the largest difference seen in the aminoglycoside and macrolide classes. However, there are large differences in the number of PNECRs per antibiotic. For example, within the quinolone class, there are 19 PNECRs that can be compared (i.e. have PNECRs in both this and in other studies). Of those 19 PNECRs, these belong to three different antibiotics (ciprofloxacin, norfloxacin, and ofloxacin), with ciprofloxacin being the most studied (n=13).

**Figure 2.**
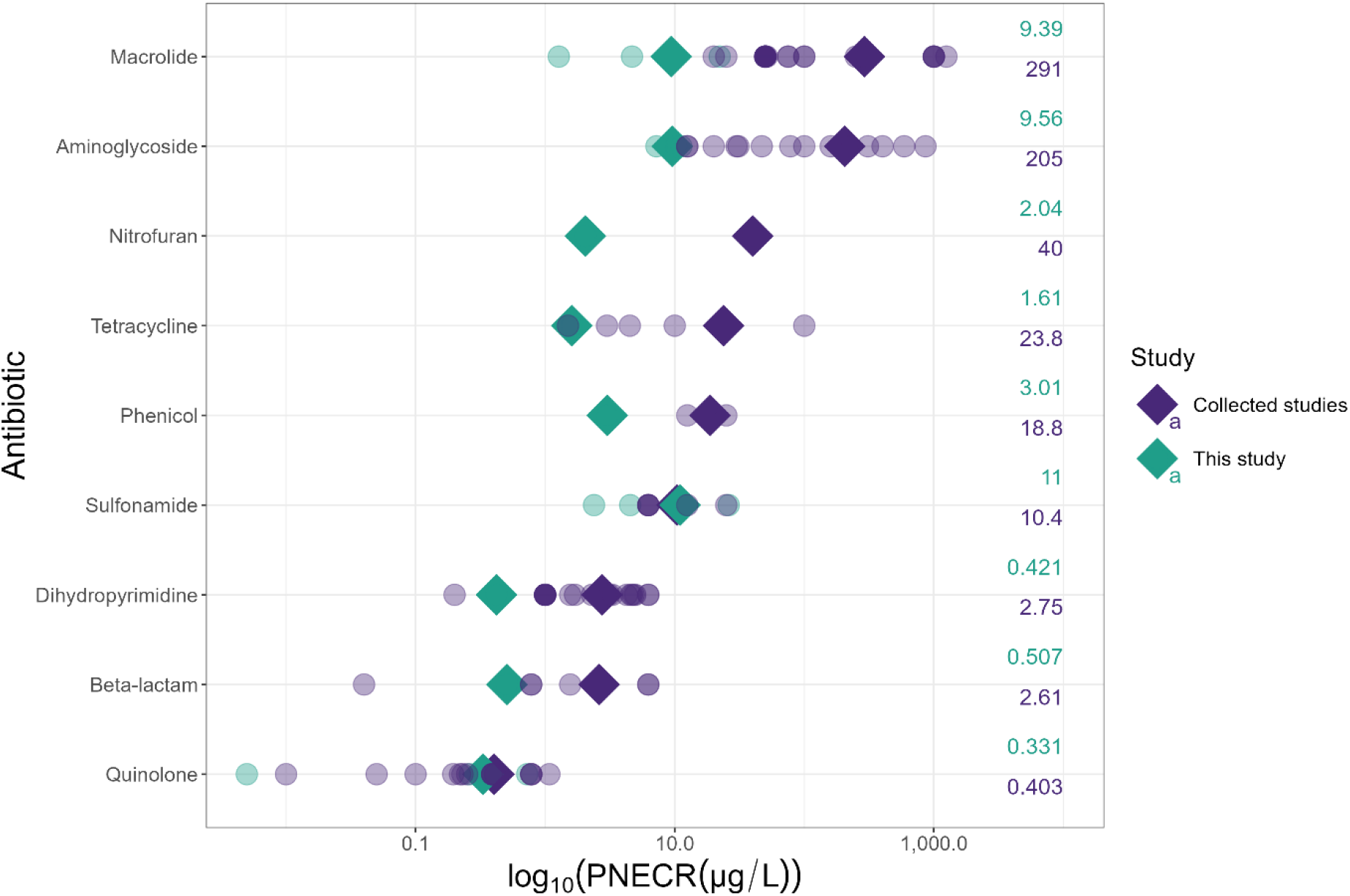
Log_10_(PNECR) comparison between those generated in this study using SELECT 2.0 and those from previous works (collected studies). Individual antibiotics are indicated by paler circles, means are indicated by bold diamonds. Labels indicate mean PNECR per class, rounded to three significant figures.

## 4 Discussion

### 4.1 SELECT 2.0 as a refined statistical method

SELECT [8] was developed as a fast and cost-effective tool to estimate PNECRs using a single experimental approach, requiring further validation and testing. We present here a statistical refinement to this previously published experimental approach to generate growth-based estimated selective concentrations within a 24-hour period and use it to generate PNECR data for 32 antibiotics across 11 classes. We propose version 2.0 represents a more statistically rigorous framework compared to original, which based PNECRs on statistically derived LOECs and NOECs. Instead, version 2.0 adopts recommended principles from ecotoxicology to model dose response relationships between antibiotic concentration and bacterial community growth, utilising all collected data, thereby reducing dependency on arbitrary, discrete test concentrations and providing continuous effect concentration estimates.

Estimation of confidence intervals around the EC1 also provides information regarding uncertainty, which is not possible with a LOEC/NOEC approach. Dose response curves can also be used to inform concentration ranges in future testing, for example, to repeat SELECT assays where EC1s lay outside the test concentration range; or to inform concentration ranges for longer-term, more complex and expensive validation experiments. This was not possible with the SELECT 1.0 approach, where if a LOEC was not found, continued iterative blind testing would be needed. Finally, the SELECT 2.0 is more flexible than the SELECT 1.0 approach. SELECT 2.0 is recommended to be used with kinetic OD readings, however, it could be used with single point OD reads (e.g. at mid-exponential phase). Therefore, the use of this approach could make data generation even more high throughput.

We used the effect concentration of 1 (EC1) for multiple reasons. Firstly, it is more protective than the EC10 recommended by the European Medicines Agency [9] and given the potential impacts of AMR in the long term [1] this seems prudent. Secondly, bacteria are fast growing (with optimal doubling time of *E. coli* around twenty minutes), meaning that population growth is significantly faster than many other ecotoxicological organisms used in chronic exposure tests where population effects may be measured over much greater time periods. Third, given the vast numbers of bacteria that will exist in natural environments, even a predicted 1% increase in resistance in a population of millions of bacteria is still a significant increase in overall numbers of resistant bacteria. Furthermore, though it has been suggested that lower effect concentrations (like EC1) are likely to be unreliable estimates due to increased confidence intervals [28], we did not find this to be the case for SELECT data (Supplementary Table 3, Supplementary Figure 1). In fact, confidence intervals were in general, smallest at EC1, likely due to the close spacing and wider range of test concentrations facilitated by the SELECT approach. We recommend that for all assays where the EC1 is calculated at the higher end of the dilution series (i.e. within the first three experimentally determined concentrations) or lies outside of the tested concentrations, the assay is repeated, with the starting concentration adjusted accordingly so the EC1 is calculated where the confidence interval is small, and therefore predictive power is increased.

Data generated by SELECT 2.0 are similar to those generated by SELECT 1.0, as evidenced by similar fold change for most tested antibiotics. Five tested antibiotics (amoxicillin, colistin, penicillin, streptomycin, and vancomycin) have SELECT 2.0 PNECRs higher than the SELECT 1.0 PNECRs. The fold changes for these differences ranged from 1.54 for colistin (SELECT 1.0 PNECR of 3.13µg/L to SELECT 2.0 PNECR of 4.80µg/L), to 10.5 for penicillin (SELECT 1.0 PNECR of 25.0µg/L to SELECT 2.0 PNECR of 263µg/L). Of those antibiotics with a lower SELECT 2.0 PNECR compared to SELECT 1.0 PNECR, we find that ceftriaxone, enrofloxacin, clarithromycin, ciprofloxacin, meropenem, chloramphenicol and nitrofurantoin have >10-fold changes in PNECR values. Furthermore, the SELECT 2.0 PNECRs are similar to many other PNECR data generated experimentally, further indicating the utility of this low-cost method to generate PNECRs. The SELECT 2.0 approach is recommended over version 1.0 moving forward owing to the open access tutorials and code and alignment with current guidelines.

Vancomycin and penicillin had the largest SELECT 2.0 PNECRs, tenfold higher than compared to the third highest PNECR. This might result from these antibiotics primarily targeting Gram-positive bacteria (of which our method does not strongly enrich for) and/or that resistance to these antibiotics is widespread and common. For example, penicillin resistant *Streptococcus pneumoniae* bloodstream infection isolates increased in prevalence in Europe by 17.5% and by 32.5% in the Africa between 2018-2023 [47]. In the UK, penicillin resistant *S. pneumoniae* isolates increased in prevalence from 1.6% to 2.7% of between 2019 and 2024 [48]. Similarly, vancomycin resistant strains are commonly found in clinical infections across the globe [49], and methicillin resistant *Staphylococcus aureus* isolates resistant to both penicillins and vancomycin are of clinical importance. Vancomycin resistance is also stable, with a prevalence of ∼20% in UK bacteraemia isolates from 2019-2024 [48]. Since we used a wastewater bacterial community in this study, it is possible that the Gram-positive taxa that were likely present (based on previous sequencing of the same community cultured under the same experimental conditions in the absence of antibiotic [50]) were already resistant to these antibiotics, reflecting this broader global picture of resistance.

The utility of a rapid, cost-effective method is the ability to generate larger, directly comparable datasets, which are currently lacking for PNECRs [17]. We have substantially increased the available data across and within antibiotic classes, enabling us to prioritise certain antibiotics and antibiotic classes for further study, risk assessments, and targeted mitigation. According to this study, the quinolone and beta-lactam antibiotics have the lowest PNECRs. By testing multiple antibiotics within classes (where possible), we were also able to identify variation of experimental putative PNECRs within multiple classes for the first time. Most classes showed little within-class PNECR variation, however for quinolones and beta-lactams within-class PNECRs varied, (from 0.00498µg/L to 1.18µg/L and 0.00003µg/L to 9.14µg/ L or 263µg/L respectively). Considering both these observations, these antibiotic classes are of greatest interest, both in terms of having the greatest selective potential and the need to study individual antibiotics. Conversely, using the approach in this study, the phenicols and aminoglycosides present little within-class variation and have higher PNECRs.

### 4.2 Future work

Below, we have discussed future work that could improve SELECT 2.0.

The PNECR data presented here are referred to as ‘putative’ as they do not have supporting data from corresponding week-long selection experiments to confirm genotypic selection for AMR. Previously, week-long exposure experiments followed by tracking of the AMR marker gene *intI1* or antibiotic class-specific resistance genes using qPCR were performed alongside SELECT assays, to verify if the SELECT PNECRs and qPCR based PNECRs were in good agreement [8]. In other words, if the growth-based, phenotypic SELECT PNECRs were suitable proxies for genotypic selection for AMR. Importantly, several antibiotics belonging to new antibiotic classes and sub-classes (e.g., glycopeptides, carbapenems) tested in this study have no genotypic data to lend further confidence to the SELECT PNECRs determined. The level of agreement between SELECT PNECRs and genotypic PNECRs may differ according to different classes, so the PNECRs reported here may not reflect genotypic selection in all cases. Future studies can perform the weeklong experiments to lend further support to the SELECT approach. The reason these were not performed in the current study is the time and resources needed to perform such assays for the number of antibiotics tested. Indeed, this is a benefit of the SELECT 2.0 method, that results can be generated in less than 24 hours, compared to several weeks at significantly greater expense. Furthermore, we argue that reduction in growth in the presence of an antibiotic by definition results in selection for resistant over sensitive strains as discussed above. Therefore, we believe that this approach is valid for generating quick, low-cost screening data that can be used to provide data for regulators or other government agencies that do not have the resources or laboratory capacity and expertise to conduct week-long experiments for all antibiotics.

The SELECT approach uses a complex bacterial community derived from sewage. This may limit the applicability of the PNECRs generated to wastewater treatment plant environments, or sewage impacted environments. There are limited studies which have compared the effects of different types of inocula [8, 12], but remarkably, results suggest PNECRs may not be substantially affected by inoculum or culturing conditions. For example, the original SELECT study evaluated the inoculum effect by comparing PNECRs from assays using influent (from two different treatment plants in the UK) and effluent samples (from one treatment plant in the UK) for four antibiotics and found the PNECR differed by a maximum of four-fold. Additionally, a recent study compared influent from the UK (Falmouth, Cornwall) and Australia (Brisbane, Queensland) and found that growth derived PNECRs were either the same or very similar across fluroquinolone, macrolide, and sulfonamide antibiotic classes [46], though these still represent only high-income countries. Future studies can test a wider variety of inocula, to better represent different environments. Different culturing conditions may be required to better represent the environments from which they are sampled, for example, lower nutrient and lower temperature conditions. Use of different media and reduced temperatures again did not have a large effect on PNECRs in the original study [8]. However, only four antibiotics were tested, so many more would be required to draw firm conclusions. Crucially, the microbial communities used as inocula should still represent the environments where the PNECRs will be used. Use of highly sensitive strains which may have poor survivability in the environment may well result in lower PNECRs, but these may then be overprotective if used in risk assessments for environments where those strains are unlikely to survive.

The default SELECT conditions (rich nutrient media, 37°C and sewage inoculum) will favour growth of certain bacterial species and reduce overall community diversity. It is currently unknown whether these conditions, which will favour human pathogens, are also more likely to favour bacteria with generally higher resistance profiles (intrinsic or acquired). Metagenome sequencing of sewage samples from the same wastewater treatment plant sampled in this study has shown that culturing under these conditions does result in species sorting, reducing overall diversity and favouring predominately growth of Enterobacterales species, though other Gram-negative and Gram-positive bacteria also persist [50–52]. Therefore, these culturing conditions could lead to an underestimation of risk. This may be a contributing factor for why certain antibiotics (e.g. narrow spectrum) tend to have higher PNECRs with this approach.

The SELECT approach is based on OD600 readings of a complex bacterial community. Whilst the inherent variability of complex communities is an advantage, because it provides a larger diversity of resistance and susceptibility profiles of different bacterial strains and species for selection to act upon, it is also a disadvantage because it will reduce overall sensitivity and potentially, reproducibility. Repeated testing, within and across laboratories, could provide data on reproducibility. It is recommended that the SELECT PNECRs are interpreted alongside other available data (see Murray et al., 2024) if attempting to define a single PNECR value for any given antibiotic.

In its current form, the SELECT approach only tests individual antibiotics, like most other methods that identify MSCs or PNECRs for antibiotics. In the environment, individual antibiotics form complex cocktails of antibiotics and potentially other co-selective compounds which may interact synergistically or even antagonistically. Though this is a limitation of virtually all studies that have investigated minimal selective concentrations and PNECRs ([37] as one of the exceptions), it could mean that selection risk is either under- or over-estimated, depending on the type of interaction effect. The SELECT approach could be modified to study mixture effects of antibiotics, other antimicrobials or co-selective compounds. However, these would need to be accompanied by corresponding weeklong selection experiments or similar, to verify that the phenotypic effect still represents genotypic selection for AMR.

## 5 Conclusions

We report the refined SELECT method (SELECT 2.0) with accompanying code and a tutorial to aid reproducibility and wider testing and validation by other groups in future studies. We found that overall, modifying the statistical approach had limited effects on PNECRs, whilst being more statistically robust and in line with policy recommendations. Given the rapid and low-cost nature of SELECT 2.0, we were able to produce the largest experimental database of experimental PNECRs generated using a single approach to our knowledge. The comparability of data enabled ranking of antibiotics and antibiotic classes according to their proposed selective potential. Overall, glycopeptides had the highest PNECRs, whereas the quinolones and dihydropyridine antibiotics tested had the lowest PNECRs. Though SELECT 2.0 still suffers from limitations that could be addressed in future studies, the accompanying open access resources will expedite future research utilising this approach. Further, the significant amount of data generated in this study will inform wider discussions around environmental risk assessment of AMR and derivation of environmental quality standards or target thresholds for antibiotic release. Reduction or prevention of selection for AMR occurring in different environments, resulting from exposure to antibiotics in anthropogenic waste streams, will only benefit human health and the environment.

## Acknowledgements

For the purpose of open access, the authors have applied a Creative Commons Attribution (CC BY) licence to any Author Accepted Manuscript version arising from this submission.

## CRediT author statement

**SK:** methodology, investigation, **AH:** data curation, methodology, investigation, validation, formal analysis, visualisation, writing – original and draft, writing – review and editing, **CL:** methodology, **WHG:** writing – review and editing, **MR:** methodology, writing – review and editing, **AB:** methodology, writing – review and editing, **AKM:** conceptualisation, methodology, validation, writing – original and draft, writing – review and editing, supervision, project administration, funding acquisition.

## Data availability statement

All code and data generated and used in analysis in this study are deposited at Zenodo DOI: https://doi.org/10.5281/zenodo.19202396. The SELECT 2.0 tutorial is available at GitHub: https://github.com/ahayyes/SELECT2.0_Tutorial.

## Abbreviations

AMR: Antimicrobial resistance
ECX: effect concentration X
ERA: environmental risk assessment
EUCAST: European Committee on Antimicrobial Susceptibility Testing
LOEC: lowest observed effect concentration
MEC: measured environmental concentration
MIC: minimal inhibitory concentration
MSC: minimal selective concentration
NOEC: no observed effect concentration
OD: optical density
PNECR: predicted no effect concentration for resistance
SELECT: SELection Endpoints in Communities of bacTeria assay

