## Supplementary File for "SELECT 2.0: Refined and open access SELection Endpoints in Communities of bacTeria (SELECT) method to determine concentrations of antibiotics that may select for antimicrobial resistance in the environment"

**Contents**

Supplementary Table 1. All antibiotics used in this study, including CAS number, product code and solvent

Supplementary Table 2. All SELECT 1.0 and 2.0 LOECs, EC1s and PNECRs generated in this study.

Supplementary Table 3. Estimates and standard error for all EC1 and EC10s for antibiotics tested in this study.

Supplementary Table 4. All PNECRs generated in this study, and those generated using data from other studies.

Supplementary Table 5. Mean PNECR per antibiotic class generated in this study, and in other collected studies. Rounded to three significant figures.

Supplementary Figure 1. Estimates and 95% confidence intervals for EC1, EC5, and EC10 for all antibiotics in this study

**Supplementary Table 1. All antibiotics used in this study, including CAS number, product code and solvent**

| **Name** | **Supplier** | **Product code** | **Cas number** | **Solvent** |
| --- | --- | --- | --- | --- |
| Amoxicillin trihydrate | Molekula | 35061441 | 61336-70-7 | DMSO |
| Ampicillin (sodium salt) | Melford | A40040_5.0 | 69-52-3 | H_2_O |
| Azithromycin | Sigma-Aldrich | PHR1088-1G | 117772-70-0 | 100% Ethanol |
| Cefotaxime sodium | Formedium | CEF0025 | 64485-93-4 | DMSO |
| Ceftiofur | Sigma-Aldrich | 34001-100MG | 80370-57-6 | DMSO |
| Ceftriaxone disodium salt hemi(heptahydrate) | Sigma-Aldrich | C5793-250MG | 104376-79-6 | H_2_O |
| Clarithromycin | Molekula | 37077446 | 81103-11-9 | 100% Acetone |
| Chloramphenicol | Sigma-Aldrich | C0378-5G | 56-75-7 | EtOH |
| Ciprofloxacin | Sigma-Aldrich | 17850-5G | 85721-33-1 | 0.1M Hydrochloric acid (use 100µL to dissolve then top up with 900uL water per mL stock) |
| Colistin sulfate | Cambridge Biosciences (Cayman Chemical Company) | CAY17584-1g (17584) | 1264-72-8 | H_2_O |
| Doripenem hydrate | Stratech Scientific (Apexbio) | A2036-APE-10MG | 364622-82-2 | H_2_O (warmed) |
| Doxycycline Hyclate | Molekula | 54888643 | 24390-14-5 | DMSO |
| Enrofloxacin | Molekula | 57134695 | 93106-60-6 | DMSO |
| Erythromycin | Acros Organics | 227330050 | 114-07-8 | 100% Ethanol |
| Florfenicol | LKT Laboratories, Inc. | F4556 | 73231-34-2 | DMSO |
| Gentamycin sulfate | Melford | G38000-5.0 | 1405-41-0 | H_2_O |
| Imipenem | Merk (Supelco.) | PHR1796-200MG | 74431-23-5 | H_2_O |
| Kanamycin sulfate | FisherScientific | BP906-5 | 25389-94-0 | H_2_O |
| Meropenem Trihydrate CRS | European Pharmcopoeis Reference Standard | Y0001252 | 119478-56-7 | H_2_O |
| Nitrofurantoin | FisherScientific | 11926821 | 67-20-9 | DMSO |
| Norfloxacin | Sigma-Aldrich | N9890-5G | 70458-96-7 | 0.5M NaOH |
| Ofloxacin | Sigma-Aldrich | 08757-10G | 82419-36-1 | DMSO |
| Oxytetracycline hydrochloride | Sigma-Aldrich | 05875-10G | 2058-46-0 | H_2_O |
| Penicillin G sodium salt | Sigma-Aldrich | 13752-1G-F | 69-57-8 | H_2_O |
| Streptomycin sulfate | Sigma-Aldrich | S9137 | 3810-74-0 | H_2_O |
| Sulfadiazine | Sigma-Aldrich | S8626-25G | 68-35-9 | DMSO |
| Sulfamethoxazole sodium salt | Molekula | 80024982 | 4563-84-2 | DMSO |
| Sulfapyridine | Molekula | 14186261 | 144-83-2 | 0.5M NaOH |
| Tetracycline hydrochloride | Sigma-Aldrich | T3383-25G | 64-75-5 | DMSO |
| Thiamphenicol | Sigma-Aldrich | T0261-1G | 15318-45-3 | DMSO |
| Trimethoprim | Sigma-Aldrich | PHR1056-16 | 738-70-5 | DMSO |
| Vanocmycin hydrochloride | Sigma-Aldrich | SBR00001-10mL | 1404-93-9 | DMSO |

**Supplementary Table 2. All SELECT 1.0 and 2.0 LOECs, EC1s and PNECRs generated in this study.**

| **Antibiotic** | **Class** | **SELECT 1.0**  **LOEC** | **SELECT 2.0**  **EC1** | **SELEC T 1.0**  **PNECR** | **SELECT 2.0**  **PNECR** |
| --- | --- | --- | --- | --- | --- |
| Amoxicillin | Beta-lactam | 62.5 | 144.553 | 3.125 | 14.4553 |
| Ampicillin | Beta-lactam | 1000 | 91.419 | 50 | 9.1419 |
| Azithromycin | Macrolide | 781.25 | 222.154 | 39.0625 | 22.2154 |
| Cefotaxime | Beta-lactam | 31.25 | 5.0675 | 1.5625 | 0.50675 |
| Ceftiofur | Beta-lactam | 62.5 | 23.5502 | 3.125 | 2.35502 |
| Ceftriaxone | Beta-lactam | 31.25 | 3e-04 | 1.5625 | 3e-05 |
| Chloramphenicol | Phenicol | 1000 | 30.116 | 50 | 3.0116 |
| Ciprofloxacin | Quinolone | 3.9063 | 0.0498 | 0.195315 | 0.00498 |
| Clarithromycin | Macrolide | 4000 | 12.744 | 200 | 1.2744 |
| Colistin | Polymyxin | 62.5 | 48.019 | 3.125 | 4.8019 |
| Doripenem | Beta-lactam | 15.625 | 3.2937 | 0.78125 | 0.32937 |
| Doxycycline | Tetracycline | 62.5 | 11.806 | 3.125 | 1.1806 |
| Enrofloxacin | Quinolone | 125 | 0.29865 | 6.25 | 0.029865 |
| Erythromycin | Macrolide | 125 | 46.805 | 6.25 | 4.6805 |
| Florfenicol | Phenicol | 937.5 | 120.89 | 46.875 | 12.089 |
| Gentamicin | Aminoglycoside | 500 | 112.311 | 25 | 11.2311 |
| Imipenem | Beta-lactam | 62.5 | 4.5051 | 3.125 | 0.45051 |
| Kanamycin | Aminoglycoside | 500 | 102.242 | 25 | 10.2242 |
| Meropenem | Beta-lactam | 125 | 1.9877 | 6.25 | 0.19877 |
| Nitrofurantoin | Nitrofuran | 500 | 20.4324 | 25 | 2.04324 |
| Norfloxacin | Quinolone | 31.25 | 7.2655 | 1.5625 | 0.72655 |
| Ofloxacin | Quinolone | 7.812 | 2.6214 | 0.3906 | 0.26214 |
| Oxytetracycline | Tetracycline | 31.25 | 12.0342 | 1.5625 | 1.20342 |
| Penicillin | Beta-lactam | 500 | 2628.27 | 25 | 262.827 |
| Streptomycin | Aminoglycoside | 31.25 | 72.252 | 1.5625 | 7.2252 |
| Sulfadiazine | Sulfonamide | 234.375 | 45.321 | 11.71875 | 4.5321 |
| Sulfamethoxazole | Sulfonamide | 250 | 23.823 | 12.5 | 2.3823 |
| Sulfapyridine | Sulfonamide | 4000 | 260.851 | 200 | 26.0851 |
| Tetracycline | Tetracycline | 250 | 16.0726 | 12.5 | 1.60726 |
| Thiamphenicol | Phenicol | 1000 | 71.799 | 50 | 7.1799 |
| Trimethoprim | Dihydropyrimidine | 31.25 | 4.2123 | 1.5625 | 0.42123 |
| Vancomycin | Glycopeptide | 8000 | 9541.4 | 400 | 954.14 |

**Supplementary Table 3. Estimates, standard error, and 95% confidence intervals for all EC1s, EC5s, and EC10s for antibiotics tested in this study.**

| **Antibiotic** | **ECX** | **Estimate** | **Standard Error** | **95% CI Lower** | **95% CI Upper** |
| --- | --- | --- | --- | --- | --- |
| Amoxicillin | EC1 | 0.144553 | 0.054278 | 0.036650923 | 0.252454 |
| Amoxicillin | EC5 | 0.341492 | 0.079658 | 0.183137967 | 0.499847 |
| Amoxicillin | EC10 | 0.50395 | 0.086973 | 0.331053092 | 0.676847 |
| Ampicillin | EC1 | 0.091419 | 0.060227 | -0.028308018 | 0.211146 |
| Ampicillin | EC5 | 0.371196 | 0.102608 | 0.167217509 | 0.575175 |
| Ampicillin | EC10 | 0.699978 | 0.161342 | 0.379240571 | 1.020715 |
| Azithromycin | EC1 | 0.222154 | 0.03864 | 0.145340924 | 0.298967 |
| Azithromycin | EC5 | 0.351275 | 0.040766 | 0.270235582 | 0.432314 |
| Azithromycin | EC10 | 0.43224 | 0.039121 | 0.354471101 | 0.51001 |
| Cefotaxime | EC1 | 0.005067 | 0.000941 | 0.003196 | 0.006939 |
| Cefotaxime | EC5 | 0.009503 | 0.001187 | 0.007143213 | 0.011864 |
| Cefotaxime | EC10 | 0.012633 | 0.001244 | 0.010159559 | 0.015106 |
| Cefriaxone | EC1 | 0.000349 | 0.000297 | -0.000241184 | 0.000938 |
| Cefriaxone | EC5 | 0.001542 | 0.000802 | -5.13E-05 | 0.003136 |
| Cefriaxone | EC10 | 0.003024 | 0.001159 | 0.000719661 | 0.005328 |
| Ceftiofur | EC1 | 0.02355 | 0.006855 | 0.009923077 | 0.037177 |
| Ceftiofur | EC5 | 0.054152 | 0.01036 | 0.033557339 | 0.074747 |
| Ceftiofur | EC10 | 0.078942 | 0.011696 | 0.055691258 | 0.102193 |
| Chloramphenicol | EC1 | 0.030116 | 0.011405 | 0.00744264 | 0.052789 |
| Chloramphenicol | EC5 | 0.112684 | 0.024977 | 0.063032064 | 0.162337 |
| Chloramphenicol | EC10 | 0.204774 | 0.032628 | 0.139912249 | 0.269636 |
| Ciprofloxacin | EC1 | 4.98E-05 | 4.67E-05 | -4.30E-05 | 0.000143 |
| Ciprofloxacin | EC5 | 0.000743 | 0.000335 | 7.75E-05 | 0.001408 |
| Ciprofloxacin | EC10 | 0.002524 | 0.000849 | 0.000836243 | 0.004211 |
| Clarithromycin | EC1 | 0.012744 | 0.008699 | -0.004548624 | 0.030037 |
| Clarithromycin | EC5 | 0.196352 | 0.11528 | -0.032818451 | 0.425522 |
| Clarithromycin | EC10 | 0.677138 | 0.421099 | -0.159979478 | 1.514256 |
| Colistin | EC1 | 0.012744 | 0.008699 | -0.004548624 | 0.030037 |
| Colistin | EC5 | 0.196352 | 0.11528 | -0.032818451 | 0.425522 |
| Colistin | EC10 | 0.677138 | 0.421099 | -0.159979478 | 1.514256 |
| Doripenem | EC1 | 0.003294 | 0.001019 | 0.001268618 | 0.005319 |
| Doripenem | EC5 | 0.005164 | 0.00111 | 0.002956881 | 0.007372 |
| Doripenem | EC10 | 0.006331 | 0.001106 | 0.004132342 | 0.008529 |
| Doxycycline | EC1 | 0.011806 | 0.002808 | 0.006223742 | 0.017388 |
| Doxycycline | EC5 | 0.036354 | 0.005495 | 0.025430814 | 0.047278 |
| Doxycycline | EC10 | 0.060487 | 0.00692 | 0.046730564 | 0.074244 |
| Enrofloxacin | EC1 | 0.000467 | 0.00019 | 8.97E-05 | 0.000845 |
| Enrofloxacin | EC5 | 0.001554 | 0.000432 | 0.000695126 | 0.002413 |
| Enrofloxacin | EC10 | 0.002677 | 0.000595 | 0.001493852 | 0.00386 |
| Erythromycin | EC1 | 0.046805 | 0.036962 | -0.026672264 | 0.120283 |
| Erythromycin | EC5 | 0.489271 | 0.396352 | -0.298651181 | 1.277194 |
| Erythromycin | EC10 | 1.415587 | 1.327799 | -1.223989497 | 4.055164 |
| Florfenicol | EC1 | 0.12089 | 0.047479 | 0.026505389 | 0.215275 |
| Florfenicol | EC5 | 0.268747 | 0.064555 | 0.140416232 | 0.397078 |
| Florfenicol | EC10 | 0.385833 | 0.067625 | 0.251398294 | 0.520268 |
| Gentamicin | EC1 | 0.112311 | 0.03084 | 0.051002332 | 0.173619 |
| Gentamicin | EC5 | 0.177516 | 0.03151 | 0.114875571 | 0.240156 |
| Gentamicin | EC10 | 0.218391 | 0.029524 | 0.159699576 | 0.277083 |
| Imipenem | EC1 | 0.004505 | 0.002231 | 7.07E-05 | 0.008939 |
| Imipenem | EC5 | 0.015039 | 0.004747 | 0.005601524 | 0.024476 |
| Imipenem | EC10 | 0.025953 | 0.006199 | 0.013629972 | 0.038275 |
| Kanamycin | EC1 | 0.102242 | 0.029328 | 0.043940229 | 0.160544 |
| Kanamycin | EC5 | 0.190752 | 0.036404 | 0.118383897 | 0.263119 |
| Kanamycin | EC10 | 0.25297 | 0.037754 | 0.17791741 | 0.328022 |
| Meropenem | EC1 | 0.001983 | 0.000518 | 0.000953231 | 0.003012 |
| Meropenem | EC5 | 0.003876 | 0.00066 | 0.002563417 | 0.005188 |
| Meropenem | EC10 | 0.005249 | 0.000689 | 0.003879056 | 0.006619 |
| Nitrofurantoin | EC1 | 0.020432 | 0.011016 | -0.001466148 | 0.042331 |
| Nitrofurantoin | EC5 | 0.106159 | 0.035188 | 0.036208016 | 0.176109 |
| Nitrofurantoin | EC10 | 0.223822 | 0.053984 | 0.116505753 | 0.331138 |
| Norfloxacin | EC1 | 0.007266 | 0.002818 | 0.001663151 | 0.012868 |
| Norfloxacin | EC5 | 0.019203 | 0.004455 | 0.010347479 | 0.028059 |
| Norfloxacin | EC10 | 0.029816 | 0.004989 | 0.019897315 | 0.039735 |
| Ofloxacin | EC1 | 0.002621 | 0.001115 | 0.000404928 | 0.004838 |
| Ofloxacin | EC5 | 0.008993 | 0.002234 | 0.004551507 | 0.013434 |
| Ofloxacin | EC10 | 0.015712 | 0.002799 | 0.010147056 | 0.021277 |
| Oxytetracycline | EC1 | 0.012034 | 0.003049 | 0.00597281 | 0.018096 |
| Oxytetracycline | EC5 | 0.031769 | 0.005252 | 0.021327726 | 0.042209 |
| Oxytetracycline | EC10 | 0.049299 | 0.006273 | 0.036828696 | 0.061768 |
| Penicillin | EC1 | 2.628275 | 0.530972 | 1.572736639 | 3.683813 |
| Penicillin | EC5 | 4.411926 | 0.582103 | 3.254743817 | 5.569108 |
| Penicillin | EC10 | 5.577759 | 0.570559 | 4.443525263 | 6.711993 |
| Streptomycin | EC1 | 0.072252 | 0.020607 | 0.031287538 | 0.113216 |
| Streptomycin | EC5 | 0.130301 | 0.02392 | 0.082749838 | 0.177852 |
| Streptomycin | EC10 | 0.170167 | 0.024051 | 0.122354833 | 0.217979 |
| Sulfadiazine | EC1 | 0.045321 | 0.01165 | 0.022161714 | 0.06848 |
| Sulfadiazine | EC5 | 0.115542 | 0.01923 | 0.077314775 | 0.153769 |
| Sulfadiazine | EC10 | 0.176491 | 0.022516 | 0.131730735 | 0.221252 |
| Sulfamethoxazole | EC1 | 0.023823 | 0.0052 | 0.013486641 | 0.034159 |
| Sulfamethoxazole | EC5 | 0.062514 | 0.009137 | 0.044349894 | 0.080678 |
| Sulfamethoxazole | EC10 | 0.096747 | 0.011119 | 0.074643283 | 0.118851 |
| Sulfapyridine | EC1 | 0.260851 | 0.088882 | 0.084159174 | 0.437543 |
| Sulfapyridine | EC5 | 0.565989 | 0.128558 | 0.310424658 | 0.821554 |
| Sulfapyridine | EC10 | 0.803699 | 0.143028 | 0.519368613 | 1.08803 |
| Tetracycline | EC1 | 0.016073 | 0.009939 | -0.003685632 | 0.035831 |
| Tetracycline | EC5 | 0.044245 | 0.016824 | 0.010800477 | 0.077689 |
| Tetracycline | EC10 | 0.069974 | 0.019486 | 0.031237378 | 0.108711 |
| Thiamphenicol | EC1 | 0.071799 | 0.023004 | 0.026068652 | 0.11753 |
| Thiamphenicol | EC5 | 0.237611 | 0.042601 | 0.152922849 | 0.322298 |
| Thiamphenicol | EC10 | 0.408451 | 0.051221 | 0.30662766 | 0.510274 |
| Trimethoprim | EC1 | 0.004212 | 0.001473 | 0.00128417 | 0.00714 |
| Trimethoprim | EC5 | 0.010013 | 0.002234 | 0.005572602 | 0.014454 |
| Trimethoprim | EC10 | 0.014818 | 0.002493 | 0.009861691 | 0.019775 |
| Vancomycin | EC1 | 9.541396 | 1.937926 | 5.688926785 | 13.39386 |
| Vancomycin | EC5 | 18.82716 | 2.385889 | 14.08416634 | 23.57015 |
| Vancomycin | EC10 | 25.60948 | 2.414956 | 20.80870533 | 30.41025 |

**Supplementary Table 4. All PNECRs generated in this study, and those generated using data from other studies.**

| **Antibiotic** | **Class** | **PNECR (µg/L)** | **Study** |
| --- | --- | --- | --- |
| Gentamicin | Aminoglycoside | 11.2311 | this |
| Gentamicin | Aminoglycoside | 20 | other |
| Gentamicin | Aminoglycoside | 862 | other |
| Gentamicin | Aminoglycoside | 12.5 | other |
| Gentamicin | Aminoglycoside | 12.5 | other |
| Kanamycin | Aminoglycoside | 10.2242 | this |
| Kanamycin | Aminoglycoside | 47 | other |
| Kanamycin | Aminoglycoside | 31 | other |
| Kanamycin | Aminoglycoside | 402 | other |
| Streptomycin | Aminoglycoside | 7.2252 | this |
| Streptomycin | Aminoglycoside | 160 | other |
| Streptomycin | Aminoglycoside | 78 | other |
| Streptomycin | Aminoglycoside | 100 | other |
| Streptomycin | Aminoglycoside | 310 | other |
| Streptomycin | Aminoglycoside | 590 | other |
| Streptomycin | Aminoglycoside | 30 | other |
| Streptomycin | Aminoglycoside | 220 | other |
| Cefotaxime | Beta-lactam | 0.50675 | this |
| Cefotaxime | Beta-lactam | 1.56 | other |
| Cefotaxime | Beta-lactam | 0.78 | other |
| Cefotaxime | Beta-lactam | 0.78 | other |
| Cefotaxime | Beta-lactam | 6.25 | other |
| Cefotaxime | Beta-lactam | 6.25 | other |
| Cefotaxime | Beta-lactam | 0.04 | other |
| Trimethoprim | Dihydropyrimidine | 0.42123 | this |
| Trimethoprim | Dihydropyrimidine | 3.3 | other |
| Trimethoprim | Dihydropyrimidine | 0.2 | other |
| Trimethoprim | Dihydropyrimidine | 2.3 | other |
| Trimethoprim | Dihydropyrimidine | 1.7 | other |
| Trimethoprim | Dihydropyrimidine | 4.96 | other |
| Trimethoprim | Dihydropyrimidine | 4.83 | other |
| Trimethoprim | Dihydropyrimidine | 4.68 | other |
| Trimethoprim | Dihydropyrimidine | 4.58 | other |
| Trimethoprim | Dihydropyrimidine | 4.27 | other |
| Trimethoprim | Dihydropyrimidine | 1 | other |
| Trimethoprim | Dihydropyrimidine | 1 | other |
| Trimethoprim | Dihydropyrimidine | 1 | other |
| Trimethoprim | Dihydropyrimidine | 1 | other |
| Trimethoprim | Dihydropyrimidine | 1 | other |
| Trimethoprim | Dihydropyrimidine | 1 | other |
| Trimethoprim | Dihydropyrimidine | 1 | other |
| Trimethoprim | Dihydropyrimidine | 6.25 | other |
| Trimethoprim | Dihydropyrimidine | 6.25 | other |
| Trimethoprim | Dihydropyrimidine | 3.13 | other |
| Trimethoprim | Dihydropyrimidine | 1.56 | other |
| Azithromycin | Macrolide | 22.2154 | this |
| Azithromycin | Macrolide | 50 | other |
| Azithromycin | Macrolide | 50 | other |
| Azithromycin | Macrolide | 50 | other |
| Azithromycin | Macrolide | 50 | other |
| Azithromycin | Macrolide | 75 | other |
| Azithromycin | Macrolide | 75 | other |
| Azithromycin | Macrolide | 25 | other |
| Azithromycin | Macrolide | 50 | other |
| Clarithromycin | Macrolide | 1.2744 | this |
| Clarithromycin | Macrolide | 250 | other |
| Clarithromycin | Macrolide | 50 | other |
| Clarithromycin | Macrolide | 1000 | other |
| Clarithromycin | Macrolide | 1000 | other |
| Clarithromycin | Macrolide | 50 | other |
| Clarithromycin | Macrolide | 50 | other |
| Erythromycin | Macrolide | 4.6805 | this |
| Erythromycin | Macrolide | 300 | other |
| Erythromycin | Macrolide | 20 | other |
| Erythromycin | Macrolide | 1250 | other |
| Erythromycin | Macrolide | 50 | other |
| Erythromycin | Macrolide | 51.41 | other |
| Erythromycin | Macrolide | 1000 | other |
| Erythromycin | Macrolide | 1000 | other |
| Erythromycin | Macrolide | 100 | other |
| Erythromycin | Macrolide | 100 | other |
| Nitrofurantoin | Nitrofuran | 2.04324 | this |
| Nitrofurantoin | Nitrofuran | 40 | other |
| Chloramphenicol | Phenicol | 3.0116 | this |
| Chloramphenicol | Phenicol | 25 | other |
| Chloramphenicol | Phenicol | 12.5 | other |
| Ciprofloxacin | Quinolone | 0.00498 | this |
| Ciprofloxacin | Quinolone | 0.25 | other |
| Ciprofloxacin | Quinolone | 0.23 | other |
| Ciprofloxacin | Quinolone | 0.22 | other |
| Ciprofloxacin | Quinolone | 0.01 | other |
| Ciprofloxacin | Quinolone | 0.78 | other |
| Ciprofloxacin | Quinolone | 0.1 | other |
| Ciprofloxacin | Quinolone | 0.05 | other |
| Ciprofloxacin | Quinolone | 1.077 | other |
| Ciprofloxacin | Quinolone | 0.7825 | other |
| Ciprofloxacin | Quinolone | 0.41 | other |
| Ciprofloxacin | Quinolone | 0.195 | other |
| Ciprofloxacin | Quinolone | 0.39 | other |
| Norfloxacin | Quinolone | 0.72655 | this |
| Norfloxacin | Quinolone | 0.39 | other |
| Norfloxacin | Quinolone | 0.39 | other |
| Ofloxacin | Quinolone | 0.26214 | this |
| Ofloxacin | Quinolone | 0.39 | other |
| Ofloxacin | Quinolone | 0.78 | other |
| Sulfadiazine | Sulfonamide | 4.5321 | this |
| Sulfadiazine | Sulfonamide | 6.25 | other |
| Sulfadiazine | Sulfonamide | 6.25 | other |
| Sulfamethoxazole | Sulfonamide | 2.3823 | this |
| Sulfamethoxazole | Sulfonamide | 6.25 | other |
| Sulfamethoxazole | Sulfonamide | 6.25 | other |
| Sulfapyridine | Sulfonamide | 26.0851 | this |
| Sulfapyridine | Sulfonamide | 25 | other |
| Sulfapyridine | Sulfonamide | 12.5 | other |
| Tetracycline | Tetracycline | 1.60726 | this |
| Tetracycline | Tetracycline | 1.5 | other |
| Tetracycline | Tetracycline | 4.5 | other |
| Tetracycline | Tetracycline | 3 | other |
| Tetracycline | Tetracycline | 100 | other |
| Tetracycline | Tetracycline | 10 | other |

**Supplementary Table 5. Mean PNECR per antibiotic class generated in this study, and in other collected studies. Rounded to three significant figures.**

| **Class** | **This study**  **mean PNECR (µg/L)** | **Other studies**  **mean PNECR (µg/L)** | **Difference in PNECRs**  **(This study – other studies)** |
| --- | --- | --- | --- |
| Aminoglycoside | 9.56 | 205 | -196 |
| Beta-lactam | 0.507 | 2.61 | -2.10 |
| Dihydropyrimidine | 0.421 | 2.75 | -2.33 |
| Macrolide | 9.39 | 291 | -282 |
| Nitrofuran | 2.04 | 40.0 | -38.0 |
| Phenicol | 3.01 | 18.8 | -15.7 |
| Quinolone | 0.331 | 0.403 | -0.0716 |
| Sulfonamide | 11.0 | 10.4 | 0.583 |
| Tetracycline | 1.61 | 23.8 | -22.2 |

**Supplementary Figure 1. Estimates and 95% confidence intervals for EC1, EC5, and EC10 for all antibiotics in this study**

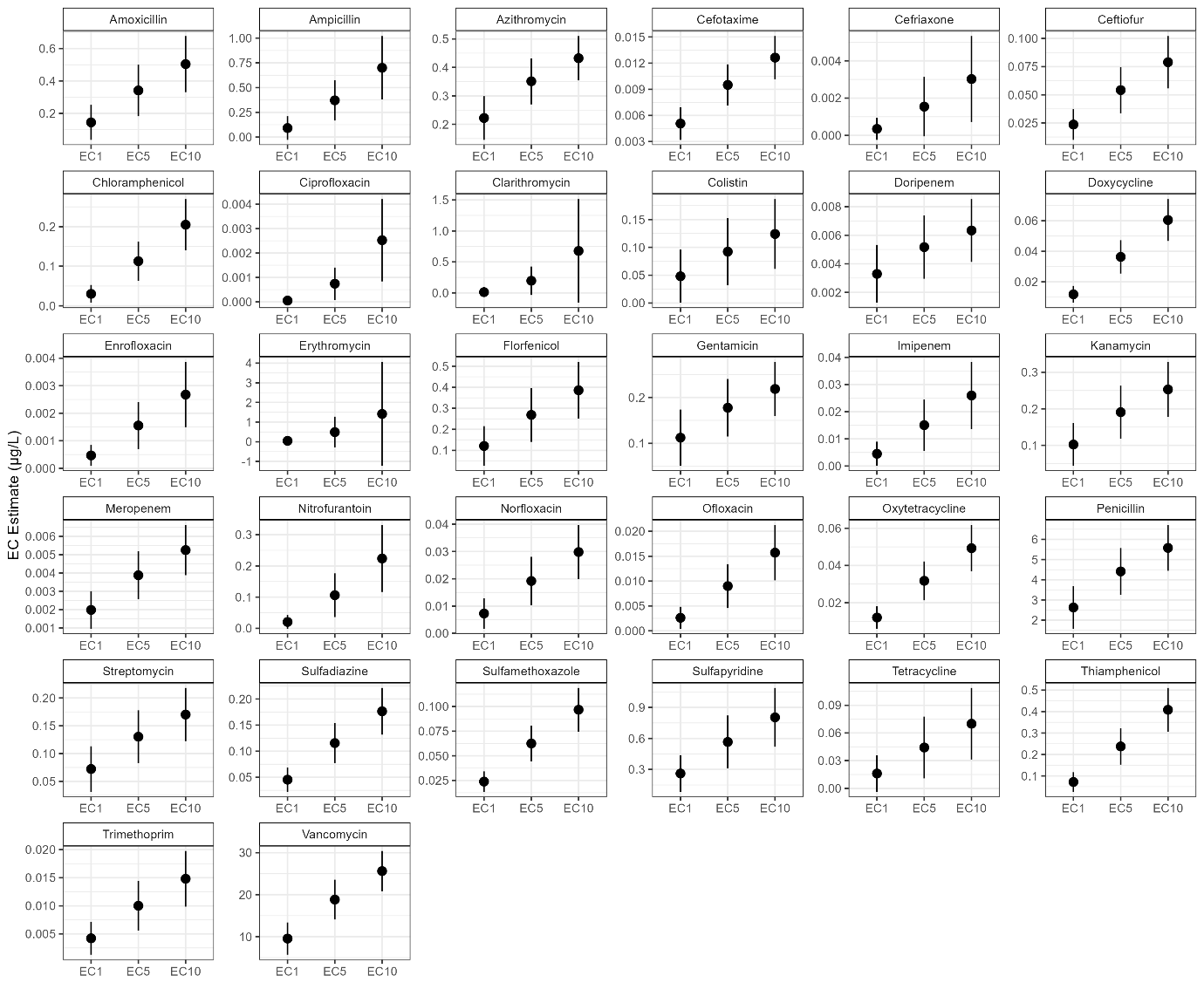
